# Core Proteome and Architecture of COPI Vesicles

**DOI:** 10.1101/254052

**Authors:** Manuel Rhiel, Bernd Hessling, Qi Gao, Andrea Hellwig, Frank Adolf, Felix T. Wieland

**Author notes:** Address correspondence to: Felix T. Wieland, Heidelberg University Biochemistry Center (BZH), Heidelberg University, Im Neuenheimer Feld 328, D-69120 Heidelberg, Germany., E-Mail, Frank Adolf, Heidelberg University Biochemistry Center (BZH), Heidelberg University, Im Neuenheimer Feld 328, D-69120 Heidelberg, Germany. Current address: MRC Laboratory of Molecular Biology, Cambridge Biomedical Campus, Francis Crick Avenue, Cambridge CB2 0QH, UK.

## Abstract

Retrieval of escaped ER-residents and intra-Golgi transport is facilitated by coat protein complex I (COPI)-coated vesicles. Their formation requires the activated small GTPase ADP-ribosylation factor (Arf) and the coat complex coatomer. Here we assess the protein composition of COPI vesicles by combining stable isotope labeling with amino acids in cell culture (SILAC) with *in vitro* reconstitution of COPI vesicles from semi-intact cells (SIC) using the minimal set of recombinant coat proteins. This approach yields an unbiased picture of the proteome of these carriers. We define a set of ~40 proteins common to COPI vesicles produced from different human as well as murine cell lines. Almost all *bona fide* COPI vesicle proteins are either ER-Golgi cycling proteins or Golgi-residents, while only a minor portion of secreted proteins was found. Moreover, we have investigated a putative role of γ- and ζ-COP as well as Arf isoforms in sorting and recruitment of specific proteins into COPI vesicles. As opposed to the related COPII system, all isoforms of coatomer and all COPI-forming isoforms of the small GTPase Arf produce COPI-coated vesicles with strikingly similar protein compositions. We present a model for the core architecture of COPI vesicles.

## Introduction

One hallmark of a eukaryotic cell is the presence of a highly complex, multiple-organelle-encompassing endomembrane system through which reaction chambers for distinct biological processes are established. Transportation of biological molecules between these compartments is largely facilitated by the trafficking of various classes of transport vesicles (Bethune and Wieland, 2018; McMahon and Mills, 2004; Pryer et al., 1992). COPII vesicles, as the first in line from the perspective of a newly synthesized protein entering the secretory pathway facilitate endoplasmic reticulum (ER)-to-Golgi transport (Barlowe et al., 1994). They are formed at specialized subdomains termed ER exit sites (ERES) by successive recruitment of the small GTPase Sar1, the heterodimer Sec23/Sec24, and the hetero-tetrameric Sec13/31 outer-coat complex (Barlowe et al., 1993; Barlowe et al., 1994; Bi et al., 2002; Matsuoka et al., 1998). The late stages of the secretory pathway are mainly served by clathrin-coated vesicles (CCVs). In this system, clathrin as the outer scaffold cooperates with various compartment-specific adaptor proteins and mostly the small GTPases of the ADP-ribosylation factor (Arf)-family or specific lipids in order to form vesicles from post-Golgi membranes (Bard and Malhotra, 2006; Bonifacino, 2004; Robinson and Pimpl, 2014; Robinson, 2015). The Golgi complex is located in between these two major trafficking systems where it serves many different functions (Wilson et al., 2011). Similar to the ER, the Golgi apparatus harbors its own vesicular transport system, namely COPI-coated vesicles (Malhotra et al., 1989; Orci et al., 1986). The small GTPase Arf, especially Arf1, plays not only a pivotal role in the formation of clathrin-coated vesicles, but also in formation of COPI vesicles (Serafini et al., 1991). Once the small GTPase is activated on the Golgi membrane by a guanosine triphosphate exchange factor (GEF), it reveals a myristoylated, amphipathic, N-terminal alpha-helix that inserts into the membrane (Antonny et al., 1997; Franco et al., 1995; Kawamoto et al., 2002; Zhao et al., 2002). Membrane-associated Arf1 in turn recruits the COPI cargo-binding and membrane scaffolding protein complex, termed coatomer, from the cytosol (Donaldson et al., 1992; Hara-Kuge et al., 1994). Stabilization of coatomer on the membrane is achieved through multiple Arf1-coatomer interactions (Eugster et al., 2000; Sun et al., 2007; Zhao et al., 1997). Additional interactions of the coat with transmembrane proteins, especially members of the p24/TMED-family, lead to the formation of a productive vesicle (Bremser et al., 1999; Cosson et al., 1998; Sohn et al., 1996). These steps of vesicle formation can be recapitulated in *in vitro* reconstitution experiments from Golgi-enriched membrane fractions, liposomes, and semi-intact cells (Adolf et al., 2013; Bremser et al., 1999; Orci et al., 1993; Spang et al., 1998).

Coatomer is a stable complex that consists of the seven subunits α-, β-, β’-, γ -, δ -, ε-, and ζCOP (Waters et al., 1991). Two of these, γ- and ζ-COP, were shown to exist as isoforms in mammals termed γ_1/2_- and ζ_1/2_-COP, respectively (Futatsumori et al., 2000). Following their initial identification, it was subsequently revealed that all of these isoforms are being incorporated into functional coatomer complexes (Wegmann et al., 2004) and can give rise to COPI vesicles *in vitro* (Sahlmuller et al., 2011). Whether the coatomer isoforms serve more diverging functions, however, remains largely elusive. In a previous study our laboratory found that γ- and ζ-COP isoforms have different preferential intracellular localizations. While γ_1_- and ζ_2_-COP were predominantly found at the cis-Golgi, γ_2_-COP displayed a more trans-Golgi localization (Moelleken et al., 2007). More recently is was revealed that the cytosolic protein Scyl1 binds to a specific class of Arfs and γ_2_-COP (Hamlin et al., 2014).

Similarly, in mammals, six isoforms of Arf have been identified (Bobak et al., 1989; Kahn et al., 1991; Price et al., 1988; Tsuchiya et al., 1991). All isoforms except for Arf2 are expressed in humans. They can be grouped into three classes based on comparison of their protein sequences and intro/exon boundaries (Kahn et al., 2006). Arf1, Arf2, and Arf3 constitute class I, Arf4 and Arf5 form class II leaving Arf6 as the sole member of class III (Kahn et al., 2006; Tsuchiya et al., 1991). Arf6 is furthermore distinguished from all other Arf family members with regard to its intracellular localization. In contrast to Arf1-5, which are recruited from the cytosol to the endomembrane system upon activation with the stable GTP analogue GTPγS and are sensitive to Brefeldin A (BFA)-treatment, Arf6 is firmly associated with the plasma membrane (Cavenagh et al., 1996). Moreover, a very thorough study by Volpicelli-Daley et al. showed a functional redundancy of class I/II Arfs when single knock-down experiments were performed, and a high specificity of individual Arfs when knock-downs targeted two Arfs at the same time (Volpicelli-Daley et al., 2005). Previous work from our lab assessed the potency of individual Arfs with respect to their ability to bind to Golgi membranes and function in COPI vesicle biogenesis (Popoff et al., 2011). It was shown that all human Arf isoforms, except for Arf6, are capable of doing both (Popoff et al., 2011). Despite this knowledge it still remains elusive, what function the different isoforms of Arf serve in COPI biogenesis at the molecular level.

Considering the differential localization of the γ- and ζ-COP isoforms (Moelleken et al., 2007) and the finding that different Golgi-tethers capture distinct populations of COPI vesicles (Malsam et al., 2005; Wong and Munro, 2014) one can posit that different isoforms of coatomer – and possibly Arf-give rise to distinct populations of COPI vesicles *in vivo*. One prediction of this concept is that vesicles formed predominantly or exclusively by one coatomer isoform or a distinct isoform of Arf would contain a specific set of cargo molecules. This would be similar to the mammalian COPII system in which coat protein isoforms are engaged in the differential sorting of transmembrane proteins into vesicles. For example all Q-SNAREs of the ER-Golgi-SNARE complex are exclusively enriched in COPII vesicles by interaction with the Sec24 isoforms Sec24C/D, whereas the R-SNARE Sec22b is specifically recognized by Sec24A/B (Adolf et al., 2016; Mancias and Goldberg, 2007, 2008). These findings have mainly emerged from studies that employed *in vitro* reconstitution systems.

Several biochemical fractionation approaches coupled to quantitative mass spectrometry (MS) analysis have been developed to globally map the localization of cellular proteins (Dunkley et al., 2004; Foster et al., 2006; Gilchrist et al., 2006). While such strategies were mostly applied for large cellular organelles, even smaller subcompartments such as transport vesicles have been objectives of such investigations (Borner et al., 2006; Gilchrist et al., 2006). Studying the protein content of vesicles has proven a useful tool to assess their biological function (Borner et al., 2006; Gilchrist et al., 2006; Takamori et al., 2006). To this end, mainly two strategies were applied. In the first setup, a fraction enriched in endogenous clathrin-coated vesicles was purified from wild type cells and compared to a corresponding fraction from clathrin heavy chain (CHC) knockdown cells (Borner et al., 2006). This experimental setup was subsequently refined by the introduction of SILAC labeling (Ong et al., 2002) and rapid mislocalization of various clathrin adaptor proteins (“knock sideways”) instead of a global CHC knockdown to investigate the role of these adaptors in greater depth (Hirst et al., 2012). The second, general approach capitalizes on classical biochemical reconstitution. Gilchrist and colleagues used a membrane fraction enriched in Golgi and cytosol to produce COPI vesicles and subsequently investigated their protein content in a label-free MS setup (Gilchrist et al., 2006).

Recently, we have combined SILAC with *in vitro* reconstitution of COPII vesicles to assess their proteome (Adolf et al., bioRxiv 253229). Here we use the same strategy to revisit the protein composition of COPI vesicles with regard to their sources and to assess the protein compositions of isotypic COPI vesicles. SILAC proteomics of the purified vesicles allowed us to i) systematically investigate the protein compositions of COPI vesicles from various cell types and ii) challenge a putative role of γ- and ζ-COP as well as Arf isoforms in the sorting of cargo molecules into these vesicles.

## Results and Discussion

### *In vitro* reconstitution of COPI vesicles for quantitative mass spectrometry

In order to study the protein content of COPI vesicles within an unbiased setup we decided to take advantage of a previously established methodology to produce such vesicles *in vitro*. Briefly, the minimal cytosolic machinery to reconstitute COPI vesicles – coatomer (CM) and Arf1 – are first expressed in Sf9 insect cells and *E.coli*, respectively. The purified recombinant proteins are then added to digitonin-permeabilized, semi-intact cells (SIC) to promote formation of vesicles from endogenous Golgi membranes. Newly formed vesicles can then be separated from their donor compartments via centrifugation (work-flow outlined in Fig. 1A).

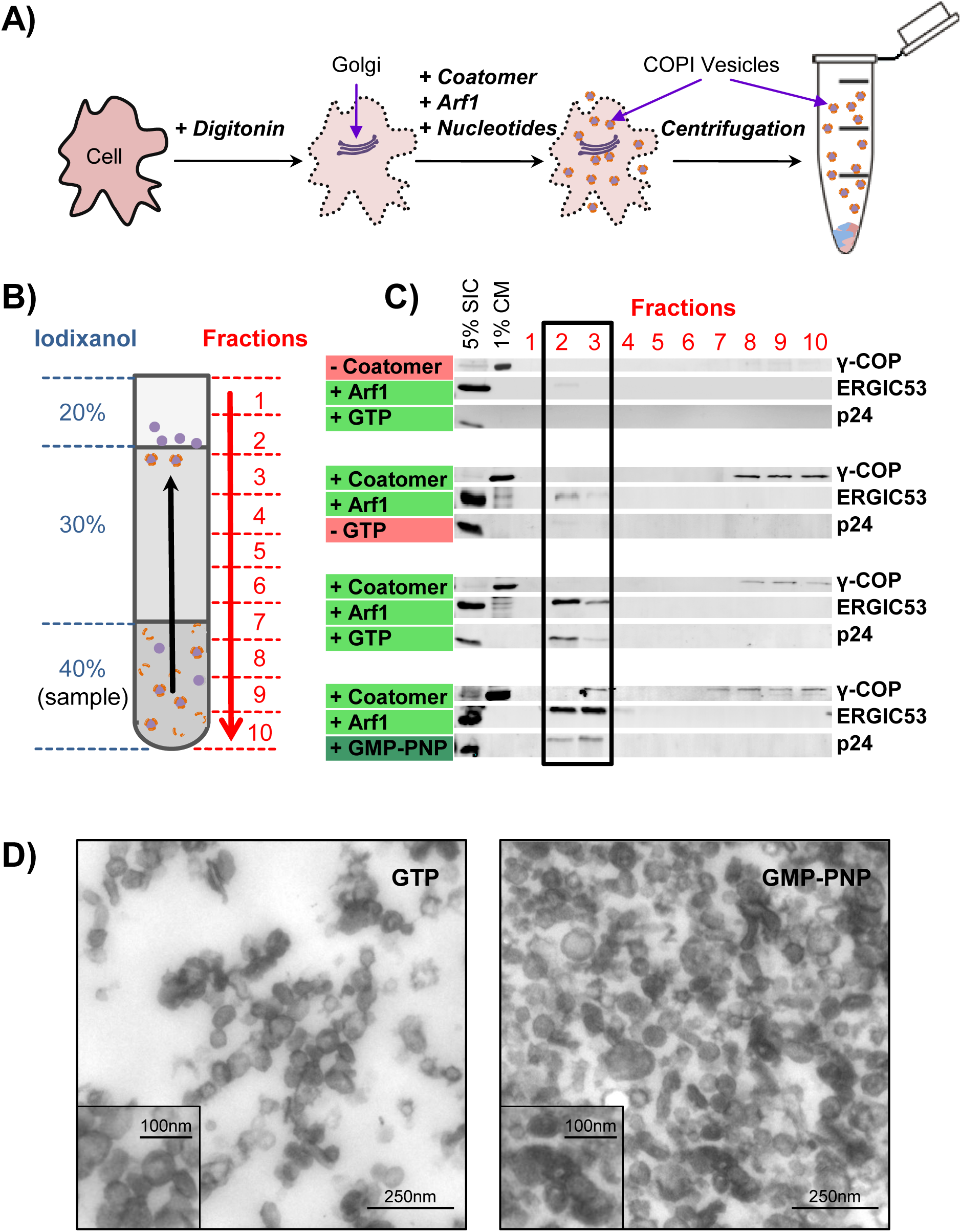
*In vitro* reconstitution and purification of COPI vesicles from semi-intact cells (SIC) A) Schematic of COPI vesicle *in vitro* reconstitution from SIC. Cells are permeabilized with digitonin and incubated with recombinant coatomer, Arf1, and guanine-nucleotide. Vesicles are separated from their donor membranes by centrifugation (see Methods section). B) Iodixanol density gradient for floatation of COPI vesicles. Schematic of the gradient used for vesicle flotation showing the different concentrations of the density matrix. The red arrow indicates fractionation from top to bottom. C) Analysis of fractions from iodixanol gradients. Ten fractions were taken from top (1) to bottom (10) and analyzed for the presence of the COPI marker proteins p24, ERGIC53 and the coatomer subunit γ-COP by western blotting. The black box highlights the fractions that contain COPI vesicles. D) Electron microscopic images of resin-embedded, COPI vesicles, which have been reconstituted *in vitro* either with GTP or GMP-PNP and then purified by floatation within an iodixanol gradient (Fractions 2+3).

However, when vesicle-containing fractions obtained with this setup were subjected to mass spectrometric analysis, we obtained only very limited and poor-quality data (data not shown). This could be explained by the vast amounts of soluble COPI coat proteins present in these samples, which hampered mass spectrometric analysis. To overcome this problem we capitalized on a recently developed density gradient for vesicle floatation with iodixanol as gradient medium (Adolf et al., bioRxiv 253229). Figure 1B shows a scheme of this gradient. Ten fractions were collected from top (fraction 1) to bottom (fraction 10) (Fig. 1B). Gradients loaded with control samples (reconstitution without coatomer or GTP) displayed only weak signals for the COPI vesicle membrane marker proteins ERGIC53 and p24 in fractions 2 and 3. When present, coat proteins (γ-COP) remained in the load fraction when present (Fig. 1C, three lower panels). In samples reconstituted with either GTP or its non-hydrolyzable analogue GMP-PNP, much stronger signals were observed for ERGIC53 and p24 in fractions 2 and 3. In samples incubated with GTP, the strongest ERGIC53 and p24 signals were detected in fraction 2, while under GMP-PNP conditions both signals, were strongest in fraction 3. Furthermore, the use of GMP-PNP instead of GTP led to co-floatation of COPI membrane marker proteins and COPI coat (here detected via γ-COP) in fraction 3 (Fig. 1C, lower panel). The shift of the strongest vesicle membrane marker signals from fraction 3 (GMP-PNP) to fraction 2 (GTP) is in agreement with a lower buoyant density expected of vesicles that have lost their coat due to GTP hydrolysis (Fig. 1C, bottom two panels).

To further analyze if fractions 2 and 3 represent vesicle-enriched samples, we performed electron microscopy (EM). Figure 1D shows images of ultrathin sections of the resin-embedded combined fractions 2 and 3 of the samples, which had been incubated with coatomer, Arf1 plus either GTP (left) or GMP-PNP (right). These fractions contained large amounts of vesicle-like structures of a size ranging from circa 60 to 100 nm. Larger structures that could occasionally be observed most likely represent membrane fragments released during the budding procedure and/or fused vesicles. While the generated with GTP seem to be without coat (Fig. 1D, left panel), many of those vesicles reconstituted with GMP-PNP displayed an electron-dense COPI coat on their surface (Fig. 1D, right panel).

### The core components of COPI vesicles from HeLa cells

The experimental setup outlined in the previous section allowed us to reconstitute and purify COPI vesicles in amounts suitable for mass spectrometric analysis. Small amounts of ERGIC53 and to a lesser degree p24 positive membranes are released from semi-intact cells during reconstitution reactions even when coatomer or GTP was omitted (Fig. 1C, upper panels). These membrane contaminants behaved similar to reconstituted COPI vesicles and they most likely have two sources. Either they stem from vesicles produced with residual coat components originating from the donor SIC or they represent unspecifically released fragments from early secretory organelle membranes. In order to subtract all contaminant-based MS-signals from those that originate from COPI vesicles reconstituted with recombinant coat proteins, we decided to combine SILAC with *in vitro* reconstitution of COPI vesicles (Adolf et al., bioRxiv 253229). As outlined in the work-flow diagram in figure 2A, cells were grown in medium containing either ‘light’ or ‘heavy’ amino acids until an incorporation of the heavy amino acids greater than 95 % was reached. From such cells, *in vitro* reconstitution reactions were performed. In the example given, a standard COPI vesicle reconstitution from heavy cells was performed in parallel to a mock-reconstitution reaction without coatomer from light cells that reflect the protein background.

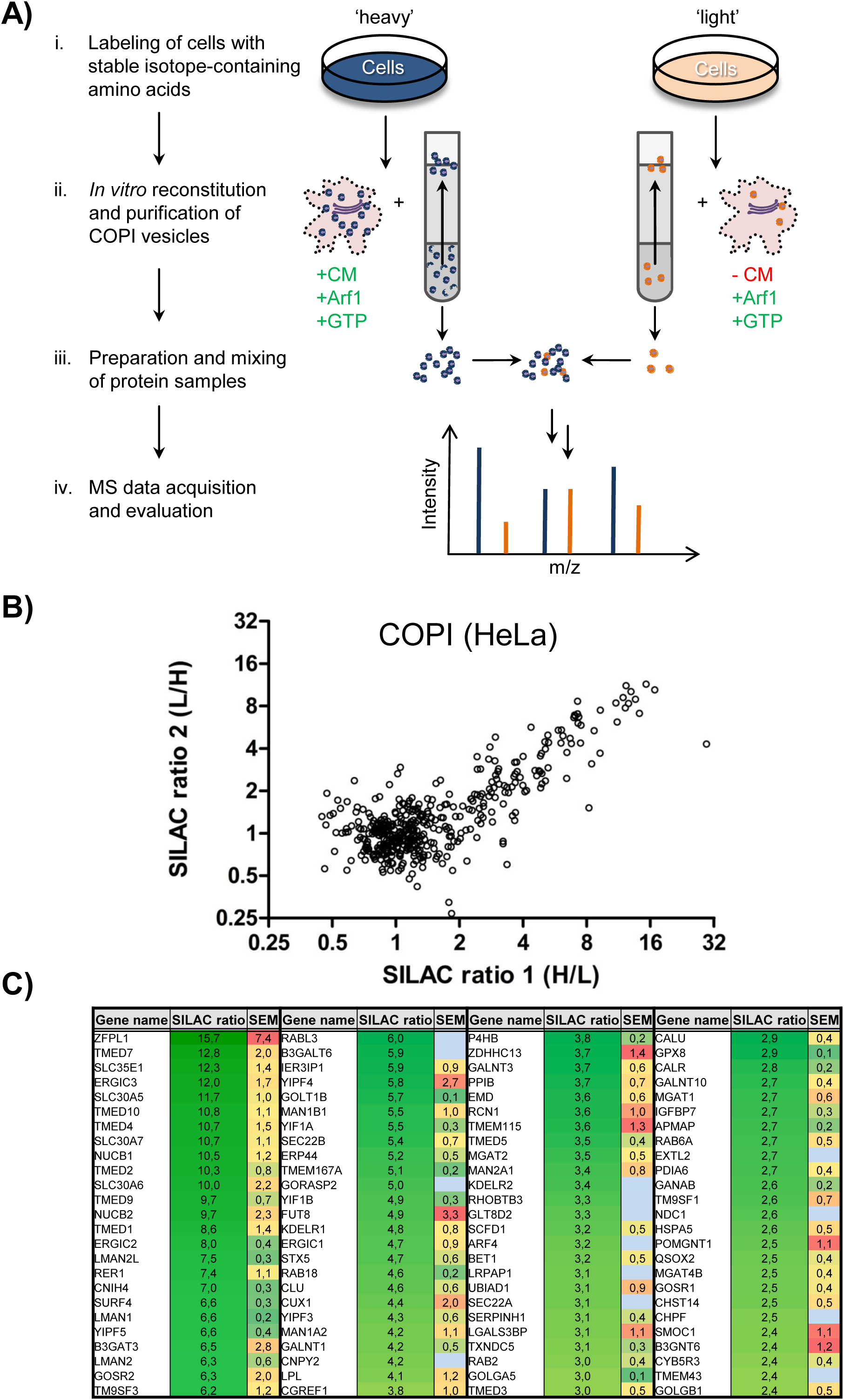
The SILAC-based core-proteome of *in vitro* reconstituted COPI vesicles. A) Work-flow of a COPI proteomic experiment. After labeling cells with heavy amino acids (i), vesicles are reconstituted and purified via floatation. In parallel, a control reaction (without CM) is performed from non-labeled cells (ii). COPI vesicle containing and control samples are isolated from the gradients, mixed 1:1 and processed (iii) to analyze both samples within the same MS run (iv) (see Methods section). B) Scatter plot representing two independent experiments as outlined in A). Experiments were performed with switched labels. C) Table of the 100 highest-scoring candidates obtained from a direct comparison of COPI vesicles with a mock reaction. Gene names, mean SILAC ratios, and standard errors of the mean (SEM) obtained from two to three experiments are shown. The few proteins with no SEM displayed were identified solely in one experiment in which the vesicle sample was produced from isotope-labeled cells.

In initial experiments not only coatomer but also Arf1 were omitted in the control reaction. Under these conditions, one adaptor protein complex-1 (AP-1) subunit was enriched fourfold over the control (data not shown). Since AP-1 is a major component of Golgi-derived CCVs, this hinted to a significant contamination of our COPI vesicle samples with AP-1/CCVs. As the formation of AP-1/CCV is also Arf1-GTP dependent (Stamnes and Rothman, 1993; Traub et al., 1993), Arf1 was included in the control experiments to allow filtering for the AP-1-dependent signal. With Arf1 included in the control, the highest SILAC ratios obtained in two data sets for an adaptor-complex subunit, AP-1 gamma, were 1.6 and 1.3, respectively (Suppl. Tab. 1). Since the coat proteins of potentially contaminating vesicle types display such low SILAC ratios, it can be expected that with this experimental setup solely vesicles of the COPI type are being recorded.

After floatation, the vesicle-containing fractions 2 and 3 from all reactions were pooled, processed by SDS-PAGE and their protein content analyzed via mass spectrometry (Fig. 2A).

To exclude possible influences of the isotopic labeling, experiments were always performed with switched labels and at least in duplicate. As an example, the SILAC ratios obtained from two independent experiments are plotted in figure 2B. Clearly, the two independent experiments yield highly correlative results. The R^2^ value obtained for the entire dataset is 0.7 and it even rises to 0.83 when the strongly diverging SILAC ratios for ZFPL1 (29.5 and 4.3) are neglected. The majority of proteins identified display SILAC ratios close to or below one, marking them as contaminants due to a close-to-equal abundance in both the heavy and the light samples. A large number of proteins, however, yielded considerably elevated SILAC ratios pinpointing them as proteins which are enriched in COPI vesicles (Fig. 2B and C). As cutoff for further analyses, a twofold enriched of candidate proteins was chosen as it is a common criterion and many known COPI proteins were found within this margin. In three “COPI reconstitution versus mock” experiments performed with HeLa cells, 102 proteins displayed a mean-enrichment of more than twofold within at least two independent experiments. An additional 20 proteins were greater than twofold-enriched in one of the measurements where the COPI membrane source was isotope-labeled, and thus represent the top COPI protein candidates. These 122 candidates (further referred to as “top 122”) constitute the COPI proteome of HeLa cells. The 100 of the top 122 proteins with the highest SILAC ratios are listed in figure 2C (complete list with the full protein names in Suppl. Tab. 1). Roughly half of the top 122 proteins were identified in all three experiments. Amongst the top 122 proteins are a large number of transport machinery proteins that cycle in the early secretory pathway (e.g. p24/TMED family proteins, LMAN1/ERGIC53, and the KDEL receptor). These proteins are known COPI vesicle constituents, e. g. members of the p24/TMED family that play a critical role in recruiting coatomer to Golgi membranes (Bremser et al., 1999; Gommel et al., 1999) and are also implicated in serving as ER-export receptor for GPI-anchored proteins (Bonnon et al., 2010). Also the well-characterized ER-Golgi cycling protein LMAN1/ERGIC53, known to directly bind to coatomer via a conserved KKXX motif at its C-terminus (Schindler et al., 1993), was enriched in the vesicle fraction. These findings underline the role of COPI carriers as shuttle between ER and Golgi.

Another group of proteins highly enriched is involved in trafficking and fusion of vesicles such as SNARE proteins (e.g. Sec22b, Stx5, and GOSR1) or Rab proteins (e.g. Rab2A, Rab6A, and Rab18). All these proteins are known to function in early steps of protein transport at the ER and/or Golgi (Dejgaard et al., 2008; Hong and Lev, 2014; Hutagalung and Novick, 2011). Furthermore, the large number of Golgi enzymes (e.g. MAN1A2, ZDHHC13, GALNT1) found in the dataset highlights the role of COPI in intra-Golgi retrograde transport. Likewise, Golgi tethers and interacting proteins are found in this COPI vesicle fraction (cf. Fig1D, e.g. ZPFL1, GOLGA5, GOLGB1).

Soluble ER-residents often carry a KDEL sequence at their C-terminus, important for their retrieval by COPI vesicles (Munro and Pelham, 1987). Accordingly several of the most abundant ER proteins (Itzhak et al., 2016) i.e. CALR, P4HB, HSPA5 (BiP), and SERPINH1 are found in the COPI proteomics dataset.

With the exception of a very few proteins (e.g. LPL, CGREF1, TGFBI, and LGALS3BP) all proteins identified can be either assigned to the ER or the Golgi complex. It is of note that NUCB1 and NUCB2 are the only soluble Golgi proteins that are found in COPI vesicles.

We further tried to identify cytosolic proteins that bind to COPI vesicles, both in their coated or uncoated state. To this end, we slightly modified our SILAC proteomics workflow (Fig. 2A) by including unlabeled or labeled cytosol to vesicle reconstitutions. Furthermore, we performed these experiments either with GTP, or the non-hydrolyzable analog GTPγS to retain the coat proteins on the vesicles (Fig. 1C). With GTPγS we noted a population of proteins clearly enriched in comparison to a reaction performed with GTP (Fig. S1B). These proteins, however, with minor exceptions, were not of cytosolic origin, but instead possess transmembrane domains and locate to the early secretory pathway, mostly the ER (e.g. translocon-associated protein subunit alpha/delta, atlastin 2/3, reticulon 1/3/4). Among the 50 proteins with the highest SILAC ratios, only two proteins are truly cytosolic (Fig.S1B, Suppl. Tab. 9 and text). Whether the cytosolic proteins are absent because they did not bind to COPI vesicles or if the experimental procedures did not allow their recovery, we cannot say. We noticed, however, that incubation of SIC with cytosol and GTPγS causes shedding of ER membranes (Fig. S1C, Suppl. Tab. 9 and text)

### Comparison of isotypic COPI vesicles

The heptameric coat building block of COPI, coatomer, comprises two subunits (γ-COP and ζ-COP) that exist as two isoforms. Coatomer with either combination of these isoforms can be purified as recombinant protein complex (Fig. 3A), and it has been shown that all can give rise to vesicles *in vitro* (Sahlmuller et al., 2011). For both γ- and ζ-COP, the two isoforms show a high level of identity: 81 % for murine γ-COP isoforms, and 73 % for ζ-COP isoforms (Fig. 3B). The most striking difference is a ~30 amino acid N-terminal extension that can be found in ζ_2_-COP (Fig. 3B).

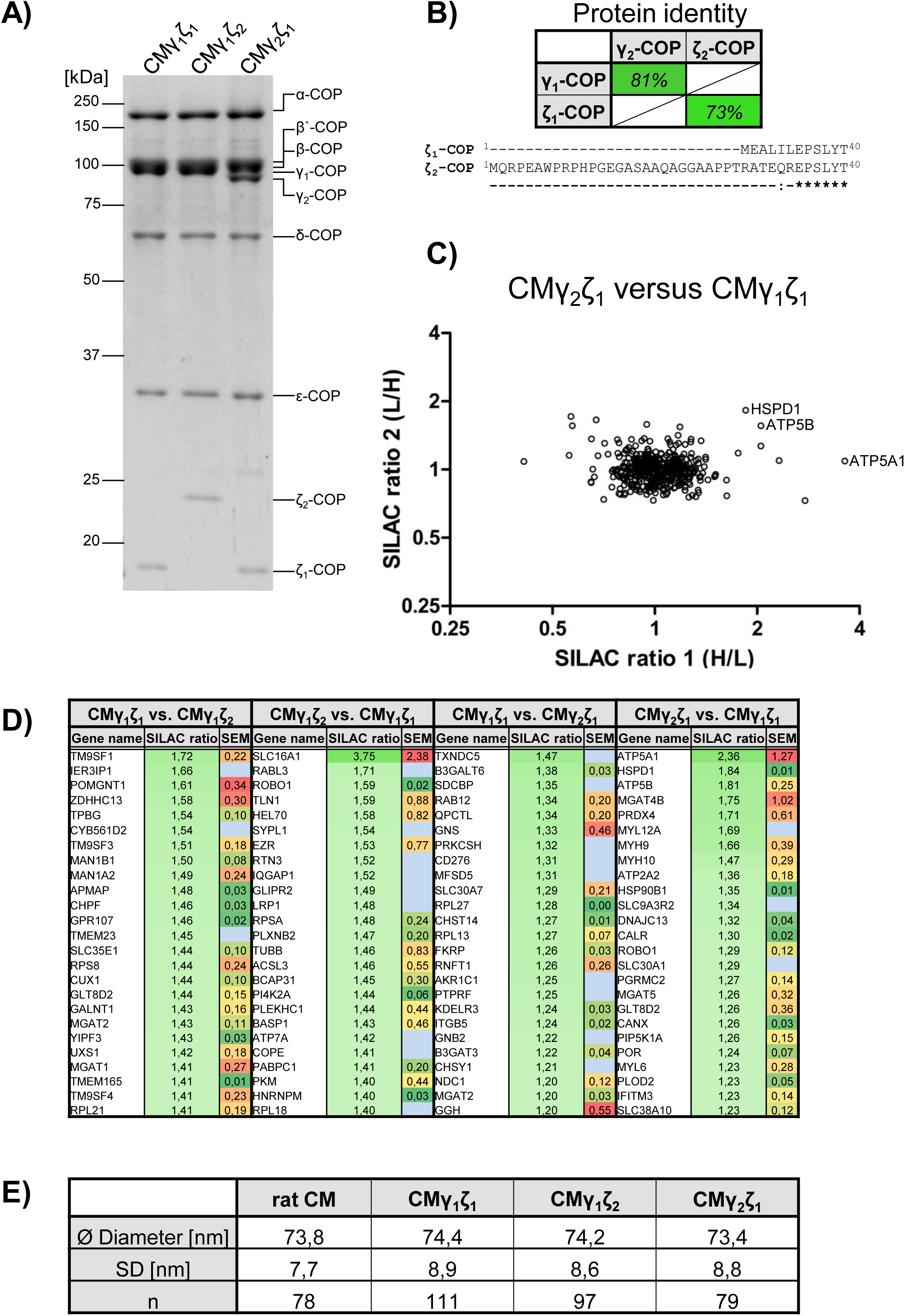
Comparison of isotypic COPI vesicles. A) Recombinant coatomer used in this study. Coomassie-stained gel of recombinant coatomer containing γ_1_-/ζ_1_-COP, γ_1_-/ζ_2_-COP, or γ_2_-/ζ_1_-COP. Subunits and apparent molecular masses are indicated. B) Representation in % of protein identity of γ- and ζ-COP isoforms (top) and alignment of the N-terminal region of ζ_1_- and ζ_2_-COP (bottom). C) Example of a scatter plot representing two independent experiments of a direct comparison of isotypic COPI vesicles. The example shows reconstitutions with γ_1_-/ζ_1_-COP-versus γ_2_-/ζ_1_-COP containing-coatomer. Experiments were performed with switched labels. Scatter plots from further comparisons of isotypic vesicles are depicted in Supplemental Material, Fig. S2A-C. D) Top 25 scoring proteins, their SILAC ratios, and standard errors of the mean (SEM) as obtained from two direct comparisons of COPI vesicles produced with different γ- and ζ-COP isoforms. Gene names, mean SILAC ratios, and SEM obtained from two experiments are shown. The few proteins with no SEM displayed were identified solely in the experiment where the respective vesicle sample was produced from isotope-labeled cells. E) Table showing the sizes of COPI vesicles generated *in vitro* with single isotypes of coatomer or an endogenous mixture of all isotypes (whole CM from rat liver).

To challenge a possible role of these isoforms in cargo selection, we decided to use the SILAC-based COPI proteomic approach as outlined in figure 2A, to directly compare the protein content of COPI vesicles made with varying coatomer isoform compositions. Figure 3C shows a scatter plot, representing two independent proteomic comparisons of vesicles produced with coatomer containing γ_1_/ζ_1_-COP (CMγ_1_ζ_1_) versus CMγ_2_ζ_1_. In contrast to a scatter plot that shows the comparison of vesicles and a mock-reaction (Fig. 2B), no relevant enrichment of proteins could be determined. In fact, the vast majority of proteins crowds at around a SILAC ratio of one.

In figure 3D the 25 proteins are listed with the highest SILAC ratios when CMγ_1_ζ_1_ is compared either with CMγ_1_ζ_2_ or CMγ_2_ζ_1_. None of the proteins showed SILAC ratios of >2 (comparison CMγ_1_ζ_1_ vs. CMγ_1_ζ_2_) or even >1.5 (comparison CMγ_1_ζ_1_ vs. CMγ_2_ζ_1_). Altogether, the isoforms 1 of both γ- and ζ- COP do not seem to select proteins in COPI vesicles different from those incorporated by the isoforms 2.

In order to further examine whether the isoforms 2 enrich proteins which are not enriched by isoforms 1, we inverted the SILAC ratios obtained in the experiments where COPI vesicles were made with CMγ_1_ ζ_1_ from heavy cells (conversion of heavy/light to light/heavy ratios). The resulting data shows, that also γ_2_- and ζ_2_-COP do not concentrate proteins in COPI vesicles different to those of their isoform counterparts (Fig. 3D). The only protein that exceeds a mean enrichment greater than twofold in the comparison of CM γ_2_ζ_1_ vs. CMγ_1_ζ_1_, ATP5A1, can be excluded as potentially isoform-specific cargo due its intra-cellular localization to mitochondria. Moreover, ATP5A1 was considerably enriched in only one of the two independent experiments (Fig. 3C and D). Other proteins that have ratios close to two, i.e. ATP5B and HSPD1, despite showing more consistent SILAC ratios (Fig. 3C and D) can also be discarded due to their localization to mitochondria.

Similarly, the comparison of CMγ_1_ζ_2_ vs. CM γ_1_ζ_1_ did not identify any isoform-specific COPI cargoes. The only protein that displayed a more than twofold enrichment was the monocarboxylate transporter 1 (SLC16A), which like ATP5A1, showed strongly divergent SILAC ratios of 6.1 and 1.4 (Fig. 3D and S2A-C).

In summary, it is more likely that the subtle changes in abundance observed for a few proteins (Fig. 3D and Suppl. Tab. 2 and Tab. 3) are the result of small differences in vesicle production and sample preparation rather than actual cargo-sorting events.

Notably, although coatomer isoforms are differentially distributed across the Golgi (Moelleken et al., 2007), Golgi enzymes that are located to specific positions within the Golgi-stack, did not show any indication of being selected by particular coatomer isotypes.

Having excluded a role of coatomer isoforms in sorting of prominent COPI cargo proteins, we decided to investigate a possible influence on another physical parameter: the size of vesicles. To this end we determined from electron microscopic images the diameter of vesicles that were reconstituted with one particular CM isoform at a time. As a control, a preparation of coatomer from rat cytosol containing all isoforms was included. EM images of reconstitutions from Golgi enriched membranes are shown in figure S3A-D. In figure S3E-F examples are given for the evaluation process. Figure S3E depicts the vesicles used for analysis marked in green, and in figure S3F just the extracted areas are shown. A summary of all measurements is given in figure 3E. The average size of COPI vesicles reconstituted under all four conditions tested varies only slightly, with diameters between 73.4 (±8.8 nm) and 74.4 (±8.9 nm), in perfect agreement with the first characterization as a (then unknown) mixture of isoforms (Orci et al., 1989).

### Comparison of Arf isoforms in COPI vesicle reconstitution

ADP-ribosylation factor (Arf) initiates the formation of COPI vesicles at the Golgi and is a stoichiometric component of their coat (Dodonova et al., 2017; Dodonova et al., 2015). Four of the five human Arf family members are capable of producing COPI vesicles (Popoff et al., 2011). This is further corroborated by the identification of Arf4 in the COPI proteome of HeLa cells as presented in the previous sections (Fig. 2C). We decided to analyze the protein content of COPI vesicles in response to usage of different recombinant Arf isoforms (Fig. 4A) for their formation. When using Arf3-5, likely due to a lower yield, less peptides and proteins were identified in COPI samples compared to Arf1. The number of proteins that are more than twofold-enriched ranges from 25 for COPI vesicles made with Arf3 to 55 for reconstitutions with Arf5 (Fig. 4B-E and Suppl. Tab. 4-6). The proteins identified in vesicles made with Arf3-5 almost entirely overlap). Twenty proteins were identified in twofold enriched in COPI vesicles reconstituted with all Arf isoforms (Fig. 4F). Among the shared proteins are 5 members of the TMED/p24 family, the SNAREs Syntaxin5 and Sec22b as well as the ER-Golgi cycling proteins ERGIC1, ERGIC2, and SURF4. Additionally, sixteen proteins are found in three vesicles types, and thirteen in COPI vesicles made with two different Arf isoforms. Most Tab. 12). In contrast, those proteins found exclusively in COPI vesicles reconstituted with a particular Arf-isoform display SILAC ratios lower than the average of the whole dataset (Suppl. Tab. 12). For example, those 20 proteins which are COPI candidates with all Arfs display a mean SILAC ratio of 8.0 in COPI vesicles made with Arf1. In contrast, those 57 proteins unique to the Arf1 candidate list show a mean ratio of 3.31 while the average ratio among all 122 candidates is 4.5 (Suppl. Tab. 12). We conclude that Arf1 is the most productive COPI-forming GTPase, however, the Arf isoforms 3-5 also produce vesicles with highly similar content.

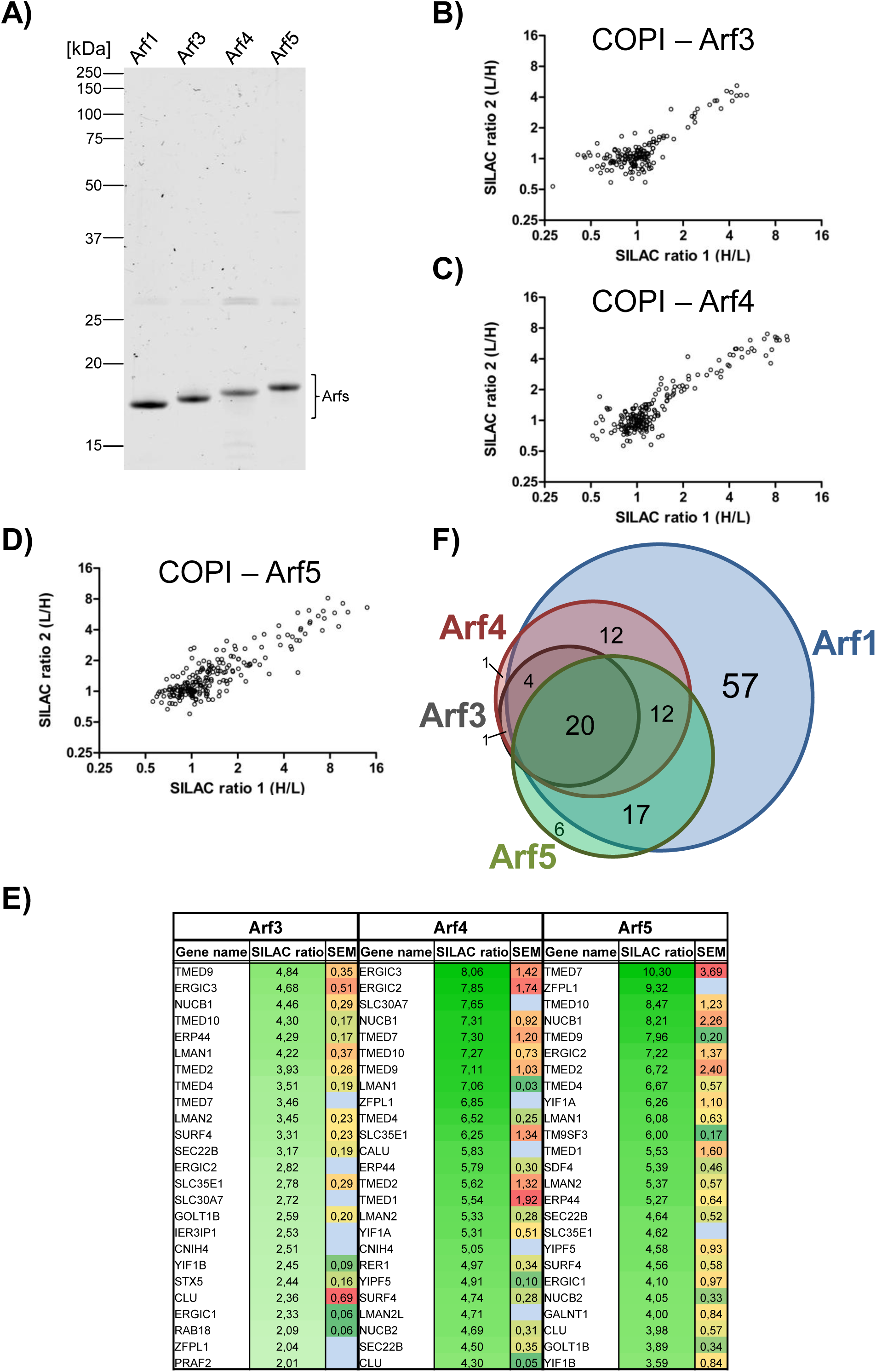
Comparison of COPI vesicles generated with various Arf isoforms. Recombinant Arf proteins used in this study. Coomassie-stained gel with recombinant Arf1, Arf3, Arf4, and Arf5. Subunits and apparent molecular masses indicated. B-D) Scatter plot representing two independent comparisons of COPI vesicles made with Arf3, Arf4, or Arf5 with mock reactions. Experiments were performed with switched labels. E) Top 25 scoring candidates, their mean SILAC ratios and standard errors of the mean (SEM) as obtained from two direct comparisons of COPI vesicles produced with the indicated Arf isoforms versus a control without coatomer. Gene names, mean SILAC ratios, and SEM obtained from two experiments are shown. The few proteins with no SEM displayed were identified solely in the experiment where the vesicle sample was produced from isotope-labeled cells. F) Venn-diagram displaying the overlap of proteins enriched &twofold in COPI vesicles made with various isoforms of Arf.

### COPI proteomics of HepG2 cells and murine macrophages

Having set up a robust assay to measure the content of COPI vesicles produced from semi-intact HeLa cells, we decided to apply the same strategy to vesicles from other cell lines in order to determine possible differences. We chose hepatocyte-like HepG2 cells as well as immortalized murine macrophages (iMΦ) as additional cell lines. As evidenced by the scatter plots shown in figures 5A and B, reproducible and robust SILAC ratios for several hundred proteins could be determined. It is of note that the SILAC ratios obtained from vesicles of HepG2 cells are on average lower than those of HeLa and iMΦ cells, with solute carrier SLC35E1 showing the highest mean ratio, 5.2. Applying the same criteria (>twofold enrichment) outlined in the previous section we were able to define a COPI proteome for HepG2 cells encompassing a total of 69 proteins. Mass spectrometric analysis of vesicle reconstitutions from iMΦ yielded 144 proteins, considerably more than we obtained from HepG2 cells. The 50 proteins with the highest SILAC ratios as found in COPI vesicles from HepG2 and iMΦ cells respectively are listed in figures 5C and D (complete lists in Tables 7 and 8 of the Supplement). A large number of proteins is found in both lists and was furthermore identified in HeLa cells (Figure 2C). A direct comparison of all proteins reveals that a shared set of 39 proteins is at least twofold enriched within the COPI proteome datasets of all three cell types (Fig. 5E and Suppl. Tab. 11). The shared 39 proteins, listed in figure 5F, account for different functional groups. In addition to e.g. fusion machinery there is a clear enrichment of early cycling proteins e.g. p24/TMED family proteins, LMAN1/ERGIC53, or Rer1 (Suppl. Tab. 11). Notably, most of these 39 conserved proteins exhibit high SILAC ratios (Figs. 2D and 5C and D). For example do these shared candidates display a mean SILAC ratio of 6.66 in HeLa cells, while candidates unique to this cell line show an average ratio of 3.45 (Suppl. Tab. 11). In addition to the 39 globally shared proteins, 18 proteins are shared only between HeLa and HepG2 cells, 19 between HeLa and iMΦ cells, and 7 proteins between HepG2 cells and iMΦ (Figs. 5E and Suppl. Tab. 11). This leaves iMΦ with 78 unique COPI proteins, whereas only 5 of the 69 COPI proteins are unique in HepG2 cells (Figs. 5E and Suppl. Tab. 11).

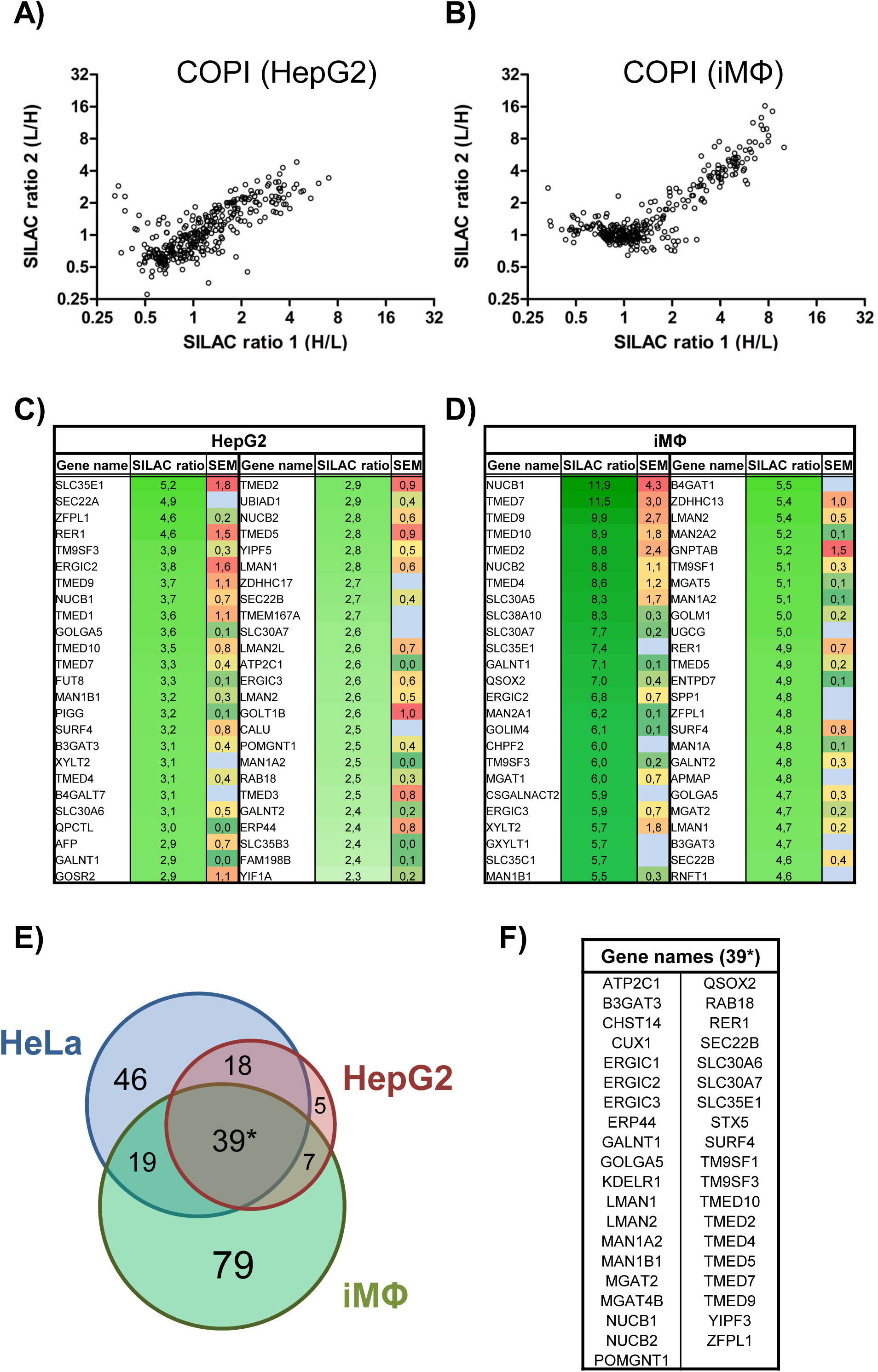
Comparison of protein compositions of COPI vesicles of various cell types. A) and B) Scatter plots representing two independent experiments that compare COPI vesicles reconstituted from HepG2 (A) or iMΦ (B) cells with a mock reaction. Experiments were performed with switched labels. C) and D) The top 50 scoring proteins within the HepG2 COPI proteome (C) or of iMΦ (D). Gene names, mean SILAC ratios, and standard errors of the mean (SEM) obtained from two experiments are shown. The few proteins with no SEM displayed were identified solely in the experiment where the vesicle sample was produced from isotope-labeled cells. E) Venn-diagram displaying the overlap of proteins that are &twofold enriched in HeLa (122 proteins), HepG2 (69 proteins), and iMΦ cells (144 proteins). F) List of the 39 proteins found >twofold enriched in all three mammalian cell lines (HeLa, HepG2, and iMΦ).

Among the conserved proteins are nonaspanins. Two of the four nonaspanin family members (TM9SF1/3) are conserved components across COPI proteomes of HeLa‐, HepG2‐ and iMΦ cells (Suppl. Tab. 11). TM9SF2 and TM9SF4, despite being identified in all three cell lines, displayed a moderate enrichment in COPI vesicles reconstituted from iMΦ, and in case of TM9SF4 also from HeLa cells. These proteins possess a C-terminal Golgi-retention motif based on the consensus-sequence KXD/E (Woo et al., 2015). Their conserved presence and the presence of many Golgi enzymes (e.g. B3GAT3, GALNT1, MAN2A1) emphasizes that COPI vesicles actively prevent the default-secretion of Golgi-residents. Moreover, enzymes involved in carbohydrate biosynthetic pathways account for the single largest group of proteins unique to iMΦ.

It is of note that overall very few soluble, secreted proteins are found with a SILAC ratio >2. Examples are the alpha-fetoprotein (AFP) found in HepG2 cells, or complement C1q subcomponent subunit beta (C1QB), while other secreted proteins, e.g. apolipoprotein E (APOE), or serotransferrin (TF) did not show any significant enrichment (Suppl. Tab. 1 and Tables 7-9). This observation is compatible with secretion via bulk-flow (Wieland et al., 1987), where active uptake into COPI vesicles is restricted to proteins for retrograde transport, while cargo for secretion is assumed to diffuse freely inside the secretory organelles and not to be concentrated at sites of vesicle production. In summary, our data highlights the presence of a strictly conserved set of proteins found in the proteomes of COPI vesicles across cell types and species. Apart from this core machinery of roughly 40 proteins (listed in Fig. 5F), various mammalian cells types seem to harbor a specific set of additional proteins.

## Conclusions

We have investigated the proteome of COPI vesicles using a novel experimental setup, combining vesicle *in vitro* reconstitution from semi-intact cells and SILAC mass spectrometry (Adolf et al., bioRxiv 253229). We could identify with high fidelity a total of 213 proteins in COPI vesicles formed from three different cell types (Suppl. Tab. 11). The largest number of proteins (144) was identified in COPI vesicles made from immortalized murine macrophages. In Hela cells 122 proteins were identified, and in HepG2 cells 69. A set of 39 proteins was present in COPI vesicles formed from all three cell lines, HeLa, HepG2, and iMΦ (Suppl. Tab. 11). These proteins, most of which cycle between the ER and the Golgi apparatus, define the basic COPI vesicle proteome and thus likely represent vesicular machinery. Many of these constituents of the COPI core proteome were found in a previous proteomic analysis of the secretory pathway, however together with several hundreds of additional proteins, which made it difficult to assign components of COPI vesicles with sufficient fidelity (Gilchrist et al., 2006).

The consistent finding of the soluble calcium binding proteins NUCB1/2 as luminal constituents of COPI vesicles, in line with their localization to the Golgi apparatus (Lin et al., 1998) and previous studies (Gilchrist et al., 2006; Rutz et al., 2009), hints towards a role of COPI vesicles in storage and release of calcium.

Another hypothetical role for a calcium binding protein as a major constituent of COPI vesicles is a function in calcium-regulated protein transport. In analogy to the pH-dependent transport of KDEL receptor clients (Wilson et al., 1993), proteins could be trafficked by NUCB1/2 (likely in concert with an additional membrane-bound protein) with their capture and release being regulated by the concentration of calcium that gradually decreases from the ER towards the trans-Golgi (Pizzo et al., 2011).

Introducing SILAC-labeled cytosol to our workflow in combination with a non-hydrolyzable GTP analog to stabilize the coat did not reveal any additional cytosolic interactors of coated COPI-vesicles (potentially except for arfaptin-1). Instead we observed that incubation of SIC with cytosol and GTPγS results in unspecific release of fragments of organelles of the early secretory pathway, mostly the ER (Suppl. material Fig. S1 and text).

The approach allowed us to study a possible influence of different isoforms of the coatomer subunits γ- and ζ-COP, as well as of the small GTPase Arf, on the content of these carriers. The protein content of COPI vesicles did not change, regardless of the various isoforms of coatomer used (Fig. 3D and Suppl. Tab. 2-3). Likewise, the protein compositions of COPI vesicles reconstituted with varying isoforms of Arf were highly similar. We noticed, however, a difference in efficiency in overall vesicle formation for differing Arfs as deduced from significantly lower numbers of peptides identified in reconstitutions with Arf3-5 as compared to Arf1. (Suppl. Tab. 12). Least proteins were identified for COPI vesicles reconstituted with Arf3. This is in line with the previous observation that Arf3 can be outcompeted from COPI vesicles by other Arf isoforms (Popoff et al., 2011).

In summary we did not find any indication that one of the different isoforms tested has a substantial influence on the content of cargo in reconstituted COPI vesicles. Moreover, we have tested a putative role of the isoforms of γ- and ζ-COP in regulating the size of COPI vesicles, but could not observe any significant difference (Fig. 3E). This leaves open the question as to the function(s) of COPI coat protein isoforms. One possibility is that the different isoforms of coatomer and Arf can transiently interact with various cytosolic or membrane proteins that are not captured in our assay. Likewise, additional cytosolic proteins may be required to modulate the content of COPI vesicles in concert with the coat protein isoforms (similar to a role attributed to GOLPH3 (Eckert et al., 2014), found at a low SILAC ratio 1.2 in vesicles made with γ_1_ ζ_1-_COPs). The only protein described to be an exclusive γ_2_-COP interactor, Scly1 (Hamlin et al., 2014), is a cytosolic protein. In our MS-analysis we did not detect Scyl1, independent of the conditions used.

Overall it seems possible that cargo capture specificity results from competition of different isoforms and is thus not observed when only one isoform is used for vesicle reconstitution. Having experimentally excluded many basic functions for coatomer isoforms in mammalian cells, a remaining attractive possibility is that isoforms of γ-COP play a pivotal role during differentiation processes in other cell types (J. Bethune, personal communication).

Figure 6 represents a schematic model of a COPI vesicle based on our proteomic study. It has been previously established that COPI vesicles form at regions of the Golgi membrane with liquid disordered phase. They contain relatively less total sphingomyelin (SM) and cholesterol than their parental Golgi membranes, whereas the molecular species SM18:0 is significantly enriched (Brugger et al., 2000) due to specific binding to the vesicular type I transmembrane protein p24 (Contreras et al., 2012). On a protein level, COPI vesicles are enriched in p24/TMED-family proteins, ER-Golgi cycling proteins/receptors (e.g. ERGIC53 and SURF4), ER-Golgi SNAREs (e.g. Stx5 and Sec22b), as well as the machinery to retrieve ER-residents (e.g. KDEL receptor and ERP44) (Fig. 6). Among the luminal proteins of COPI vesicles, NUCB1 and NUCB2 are highly abundant, present at a much higher concentration than KDEL-bearing ER-residents. Rather, these Ca-binding proteins occur in amounts about stoichiometric to the COPI membrane machinery proteins (Rutz et al., 2009). They likely interact with the luminal parts of membrane proteins to facilitate their uptake into vesicles (Fig. 6, indicated by black arrows). Whether secreted cargo can be found in COPI vesicles cannot be deduced from our proteomic study, possibly due to the generally low abundance of such proteins, as expected for cargo that undergoes non-signaled bulk uptake.

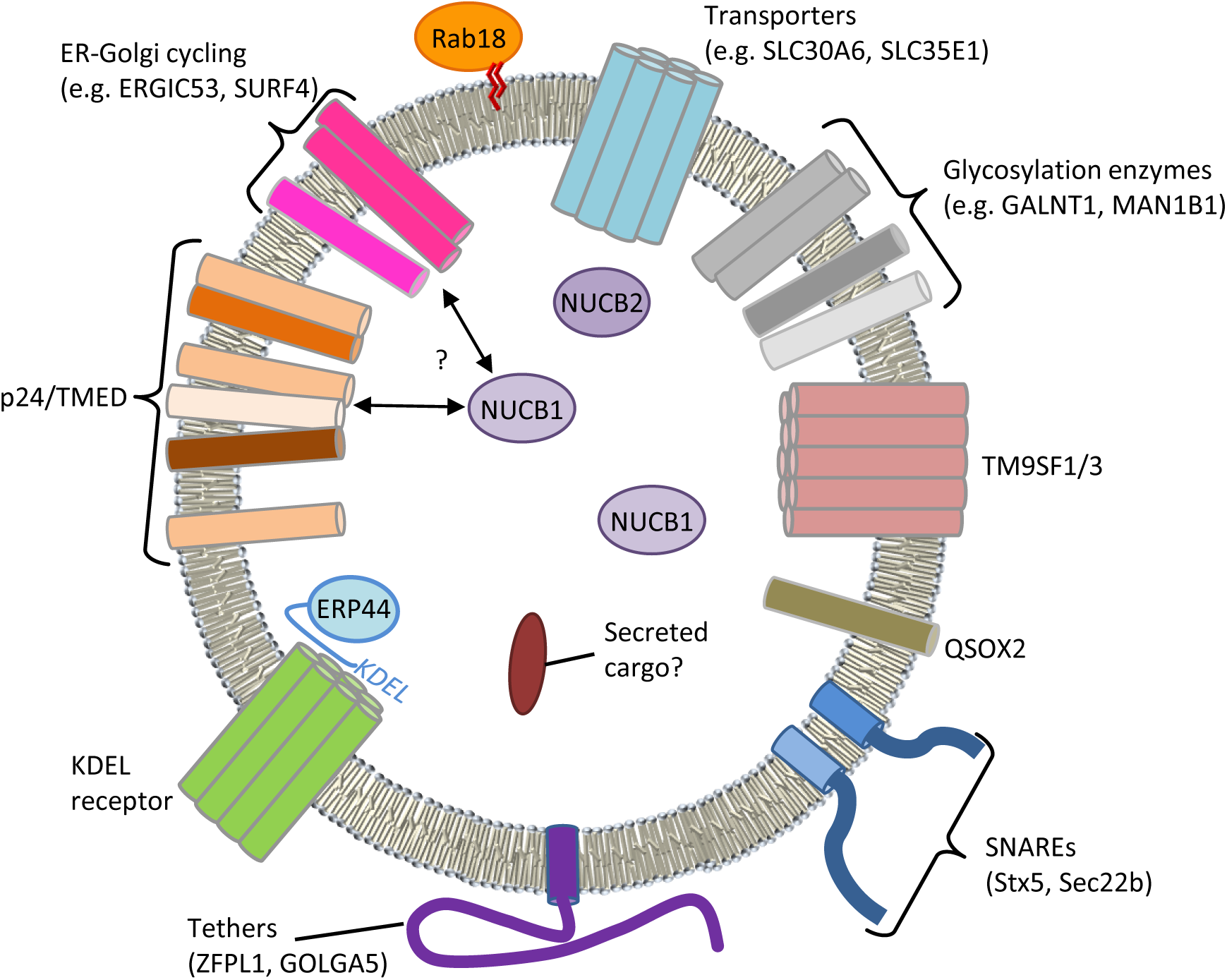
Model of a COP vesicle with its core components. Schematic representation of a COPI vesicle with its core proteome based on our SILAC proteomics data (Fig. 5F). For description and discussion please refer to the Conclusion section of the main text.

Apart from the aforementioned proteins, COPI vesicles carry factors for vesicle tethering (e.g. ZFPL1 and GOLGA5) and Golgi-resident transporters for small molecules across membranes (e.g. SLC30A6 and SLC35E1). As expected, Golgi-resident enzymes were found enriched in the vesicles that serve glycosylation (e.g. GALNT1 and MAN1B1). Moreover, proteins that contribute to the formation of disulfide bonds (QSOX2) or have a less defined function (TM9SF1 and TM9SF3) are among the constituents that define the core of an intra-Golgi and Golgi-to-ER carrier (Fig. 6 and 5F).

On the cytoplasmic side, besides the various established binding partners of membrane attached coatomer, very few proteins seem to stably interact with COPI vesicles (also see previous sections). Proteins of the Rab family were the only cytosolic proteins identified under multiple conditions. Here, Rab18 was the most persistently identified family member (Fig. 6 and 5F).

Taken together, our results define the core proteome of COPI vesicles and reveal that the various isoforms of the COPI coat do not reflect functions in differential uptake of cargo (in line with an un-signaled uptake of cargo), and leave open a possibility that isoforms serve specific purposes during steps in development.

The presence of Ca‐binding proteins in the vesicular lumen at concentrations stoichiometric to major membrane proteins (Rutz et al., 2009) hints at functions of NUCB1/2 in the molecular mechanism of cargo uptake or release in a pathway along a luminal calcium gradient, characteristic of the early secretory pathway (Pizzo et al., 2011).

## Materials and Methods

### Antibodies

First antibodies used in this study: anti-calnexin (ab75801, Abcam, UK); anti-ERGIC53 (sc-365158, Santa Cruz Biotechnology, USA); anti-GM130 (610822, BD Biosciences, USA); anti-γR-COP (Pavel et al., 1998); anti-p24 (Gommel et al., 1999). Secondary antibodies used for western blot analysis: goat anti-mouse IgG Alexa Fluor 680 conjugated (A-21058, Thermo Fisher Scientific, USA); goat anti-rabbit AlexaFluor 680 conjugated (A-21076, Thermo Fisher Scientific, USA).

### Protein expression and purification

Recombinant, myristoylated hArf paralogues (Arf1-Arf5) were expressed and purified based on previously established protocols (Popoff et al., 2011). Briefly, bacterial expression pellets usually originating from 4 liter of bacterial cultures were resuspended in 50 ml lysis buffer (50 mM Tris‐HCl pH 8.0, 1 mM MgCl_2_, 1 mM DTT, 1 mM GDP, 1 tablet cOmplete™ EDTA-free Protease Inhibitor Cocktail [Roche, Switzerland]). Cells were lysed via 5 runs through a Microfluidizer^®^ (Microfluidics, USA). The lysate was cleared of debris via centrifugation (100.000×g, 4°C, 1 h, TFT55-38 rotor [Kontron Instruments, Germany]) and Arf proteins were precipitated with ammonium sulfate at a final concentration of 40% over a course of 1.5 h. The precipitate was harvested (10.000×g, 4°C, 30 min, SLC-1500 rotor [Sorvall, USA]), resuspended in lysis buffer, and the Arf isoforms were further purified via a run over a Superdex75 (16/60) column (GE Healthcare, USA) equilibrated in storage buffer (25 mM HEPES pH 7.4 [KOH], 200 mM KCl, 5 mM MgCl_2_, 1 mM DTT, 1 mM, 10 %(w/v) glycerol). Recombinant murine coatomer containing different γ- and ζ-COP isoforms was expressed in Sf9 insect cells. The heptameric complexes were purified via a One‐STreP-Tag C-terminal of the α-COP subunit with *Strep*-Tactin^®^ Sepharose^®^ (IBA, Germany).

### Cultivation of Cells

HeLa, HepG2 and iMΦ cells were grown at 37°C with 5 % CO_2_ in DMEM medium containing either Lys-8/Arg-10 (heavy) or Lys-0/Arg-0 (light), supplemented with 10 % fetal bovine serum (SILAC-Lys8-Arg10-Kit, Silantes, Germany).

### *In vitro* reconstitution of COPI vesicles

Semi-intact cells for COPI reconstitution were prepared as previously described (Mancias and Goldberg, 2007). *In vitro* formation of COPI vesicles was essentially carried out as described by Adolf et al. (2013). Modifications for a standard budding reaction were the use of 200 µg instead of 100 µg of SIC in a volume of 200 µl together with 4 µg Arf and 10 µg OST-coatomer. Reconstitution of COPI vesicles from rat liver Golgi was performed as described by Popoff et al. (2011).

### Preparation of HeLa cell cytosol

Cells, nearly confluent, from 4-6 dishes (Ø 15 cm) were trypsinized and resuspended in PBS supplemented with trypsin inhibitor. Cells were washed once with assay buffer (25 mM HEPES pH 7.2 [KOH], 150 mM KOAc, 2 mM MgOAc) and resupended in a small volume (~1 ml) of assay buffer. Lysis was achieved through nitrogen cavitation (800 psi, 30 min, on ice) using a 4639 cell disruption vessel (Parr Instruments, USA). The soluble cytosolic fraction was cleared from debris via centrifugation at 100.000×g within a TLA45/55 rotor (Beckman Coulter, USA) at 4°C for 1 h.

### Purification of COPI vesicles via floatation within an iodixanol gradient and MS sample preparation

COPI vesicle samples reconstituted as outlined above were adjusted to 40 % of iodixanol (Sigma-Aldrich, USA) in a final volume of 700 µl and subsequently overlayed by first 1200 µl of 30 % iodixanol solution and finally 400 µl of 20 % iodixanol in assay buffer (25 mM HEPES pH 7.2 [KOH], 150 mM KOAc, 2 mM MgOAc). The density gradients were centrifuged for 13-15 h at 250.000×g in an SW60-Ti rotor (Beckman Coulter, USA). The top 200 µl of the gradient were discarded and a 500 µl vesicle-containing fraction was isolated. The vesicles were subsequently harvested by diluting the fraction 1:3 in assay buffer and subsequent centrifugation at 100.000×g in a TLA45/55 rotor (Beckman Coulter, USA) for 2 h at 4°C. The supernatant was again discarded and the samples dissolved in SDS sample buffer through boiling at 95°C for 10 min. For a single SILAC experiment, six budding‐ and control-reactions were performed in parallel from either heavy or light cells. The samples were mixed in a 1:1 ratio and briefly run (approximately 1 cm) into a 10 % Tris-Glycine gel (Thermo Fisher Scientific, USA), stained with Roti®-Blue colloidal coomassie (Roth GmbH, Germany) and further processed for mass spectrometric analysis.

### Electron microscopy of reconstituted COPI vesicles

For electron microscopic investigation, COPI vesicles, which have been reconstituted and purified as outlined above from SIC, were resin embedded as described by Adolf et al. (2013). Briefly, the yield from three gradients per sample was pooled, sequentially harvested at 100.000×g in a TLA45/55 rotor (Beckman Coulter, USA) for 1 h and further processed. COPI vesicles, which have been generated from rat liver Golgi, were negatively stained.

### Vesicle size determination

COPI vesicle in electron microscopic images show a roughly circular shape. The area encircled by the membrane has inhomogeneous pixel intensities. Due to the significant noise and low contrast of the images, common circle detection-based and pixel classification-based segmentation methods are inadequate. Since the membrane profiles were consistent, we employed the segmentation method of Dimopoulos et al. (2014), which exploits the membrane patterns and can achieve an optimal detection of object boundaries(Dimopoulos et al., 2014).

In particular, we used a two-step semi-automatic segmentation scheme: i) positions of vesicles were manually localized to further use them for initialization or as seeds for the following segmentation step. Broken vesicles and similar structures were excluded. ii) The method described by Dimopoulos et al. (2014) was applied to segment the images based on membrane profiles obtained from a few vesicles as examples. The segmented vesicles with a low score of membrane profile were discarded to guarantee an accurate segmentation. The vesicle area was finally computed using the number of pixels in the segmented vesicle and the pixel size information.

### Mass spectrometry and data analysis

Gel pieces were reduced with DTT, alkylated with iodoacetamide and digested with trypsin using the DigestPro MS platform (Intavis AG, Germany) following the protocol described by Shevchenko et al. (Shevchenko et al., 2006).

Peptides were analyzed by liquid chromatography–mass spectrometry (LCMS) using an UltiMate 3000 LC (Thermo Scientific, USA) coupled to either an Orbitrap Elite or a Q-Exactive mass spectrometer (Thermo Scientific, USA). Peptides analyzed by the Orbitrap Elite were loaded on a C18 Acclaim PepMap100 trap-column (Thermo Fisher Scientific, USA) with a flow rate of 30 µl/min 0.1 % TFA. Peptides were eluted an d separated on an C18 Acclaim PepMap RSLC analytical column (75 µm × 2 50 mm) with a flow rate of 300 nl/min in a 2 h gradient of 3 % buffer A (0.1 % formic acid, 1 % acetonitrile) to 40 % buffer B (0.1 % formic acid, 90 % acetonitrile). MS data were acquired with an automatic switch between a full scan and up to 30 data-dependent MS/MS scans.

Peptides analyzed on the Q-Exactive were directly injected to an analytical column (75 µm × 300 mm), which was self-packed with 3 µm Reprosil Pur-AQ C18 material (Dr. Maisch HPLC GmbH, Germany) and separated using the same gradient as described before. MS data were acquired with an automatic switch between a full scan and up to 15 data-dependent MS/MS scans.

Data analysis was carried out with MaxQuant version 1.5.3.8 (Cox and Mann, 2008) using standard settings for each instrument type and searched against a human or mouse specific database extracted from UniProt (UniProt Consortium). Carbamidomethylation of cysteine was specified as fixed modification; oxidation of methionine, deamidation of asparagine or glutamine and acetylation of protein N-termini was set as variable modification. ‘Requantify’ as well as ‘Match Between Runs’ options were both enabled.

Results were filtered for a 1 % false discovery rate (FDR) on peptide spectrum match (PSM) and protein level. MaxQuant output files were further processed and filtered using self-compiled R-scripts and Excel (Microsoft, USA).

## Author contributions

MR performed all reconstitutions with recombinant proteins and purification of SIC derived vesicles. BH performed MS experiments, QG analyzed EM-images for vesicle size determinations, AH performed electron microscopy experiments. FA and FW supervised coworkers, and MR and FW wrote the manuscript. All authors revised the manuscript.

## Acknowledgements

We wish to thank all members of the Wieland lab for helpful comments, fruitful discussions, and their support. We would further like to thank Eicke Latz (Institute of Innate Immunity, Bonn) for kindly providing immortalized macrophages and Julien Béthune for critical reading of the manuscript. We are grateful to Hilmar Bading (Department of Neurobiology and Interdisciplinary Center for Neurosciences, Heidelberg) for providing the opportunity to carry out the electron microscopy work in his laboratory. This work was supported by the German Research Council, SFB 638, A10 and DFG-Einzelprojekt Wi-654/12-1.

## Supplementary Material

### Proteomics Data

Tables T1-T12 contain comprehensive overviews of the mass spectrometric data that is referred to in the text. The identified proteins, their corresponding gene names and SILAC ratios are given in T1-T8. Table 11 shows the comparison of the top scoring proteins from HeLa (T1), HepG2 (T7), and iMΦ (T8). A comparison of the COPI proteomes of vesicles made with different isoforms of Arf (T1, T4-6) is shown in table 12. Detailed information on all peptides and proteins identified in this study can be found in T13-T16.

### Control experiments to assess an influence of non-hydrolyzable GTP analogs on the release of material from SIC

In experiments to analyze possible COPI interactors present in cytosol, the non-hydrolyzable GTP analog GTPγS was used in order to stabilize the vesicular coat.

Under these conditions, we detected many membrane proteins of the ER released from SIC, as compared to COPI vesicle formation with GTP. Therefore we decided to probe the cytosol used for the presence of the highly abundant ER marker calnexin and the ERGIC/cis-Golgi proteins ERGIC53 and p24. None of the proteins could be detected (Fig. S1A). Thus we excluded the cytosol as source of the membrane proteins released with GTPγS. Expectedly, soluble cytosolic proteins (Arf1, ε-COP) as well also soluble ER-content (BiP/GRP78) partitioned with the cytosolic fraction (Fig. S1A). Hence, we tested whether the use of GTPγS alone would lead to the release of early secretory pathway membrane proteins. In a reconstitution reaction with GTP γS in the presence of cytosol, but without recombinant coatomer, we again detected a large number of ER proteins. Of the 38 most-enriched proteins, 33 are also found among the 70 proteins with the highest SILAC-ratios in the sample with coatomer (Fig. S1B and C and Suppl. Tab. 10). We conclude that incubation of semi-intact cells with GTPγS and cytosol causes partial fragmentation of the ER.

## Figure Legends

**Fig. S1:**
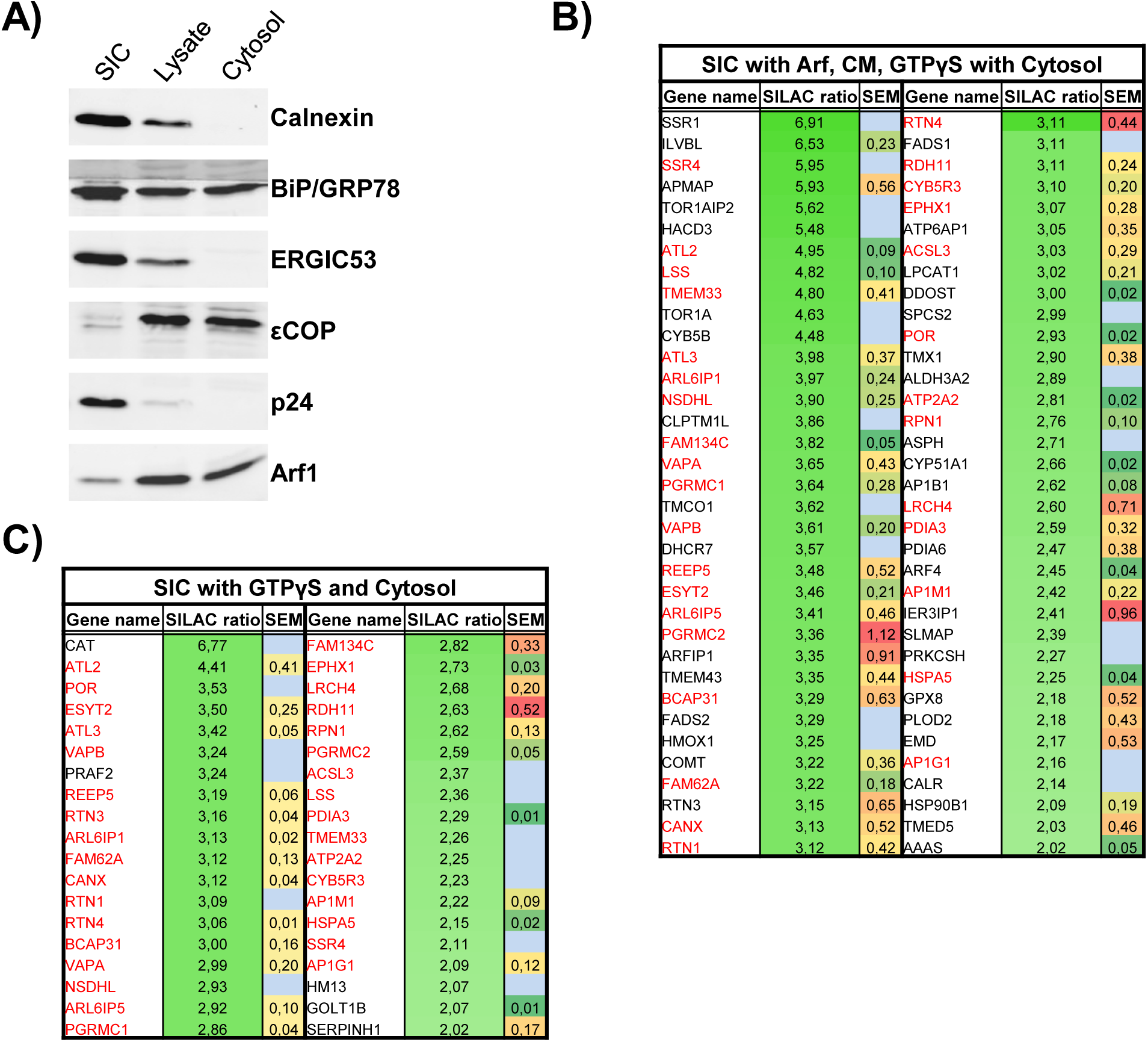
Comparison of material released from SIC in the presence of GTPγS or GTP. A) Western blot analysis of semi-intact HeLa cells (SIC), HeLa cell lysate and cytosol for the presence of the proteins indicated. B) The top72 scoring proteins (SILAC ratios of >2) from a comparison of COPI vesicles reconstituted with GTPγS or GTP in the presence of cytosol. Gene names, mean SILAC ratios, and standard errors of the mean (SEM) obtained from two independent experiments are shown. The few proteins with no SEM displayed were identified solely in the experiment where the GTPγS vesicle sample was produced from isotope-labeled cells. Gene names of proteins also found in C) are red. C) Proteins scoring SILAC ratios of &2 from samples released from SIC by cytosol, Arf1, and GTPγS or GTP. Gene names, mean SILAC ratios, and SEM as obtained from two independent experiments are shown. The few proteins with no SEM displayed were identified solely in one experiment where the GTPγS sample was produced from isotope-labeled cells. Gene names of proteins also found in B) are red.

**Fig. S2:**
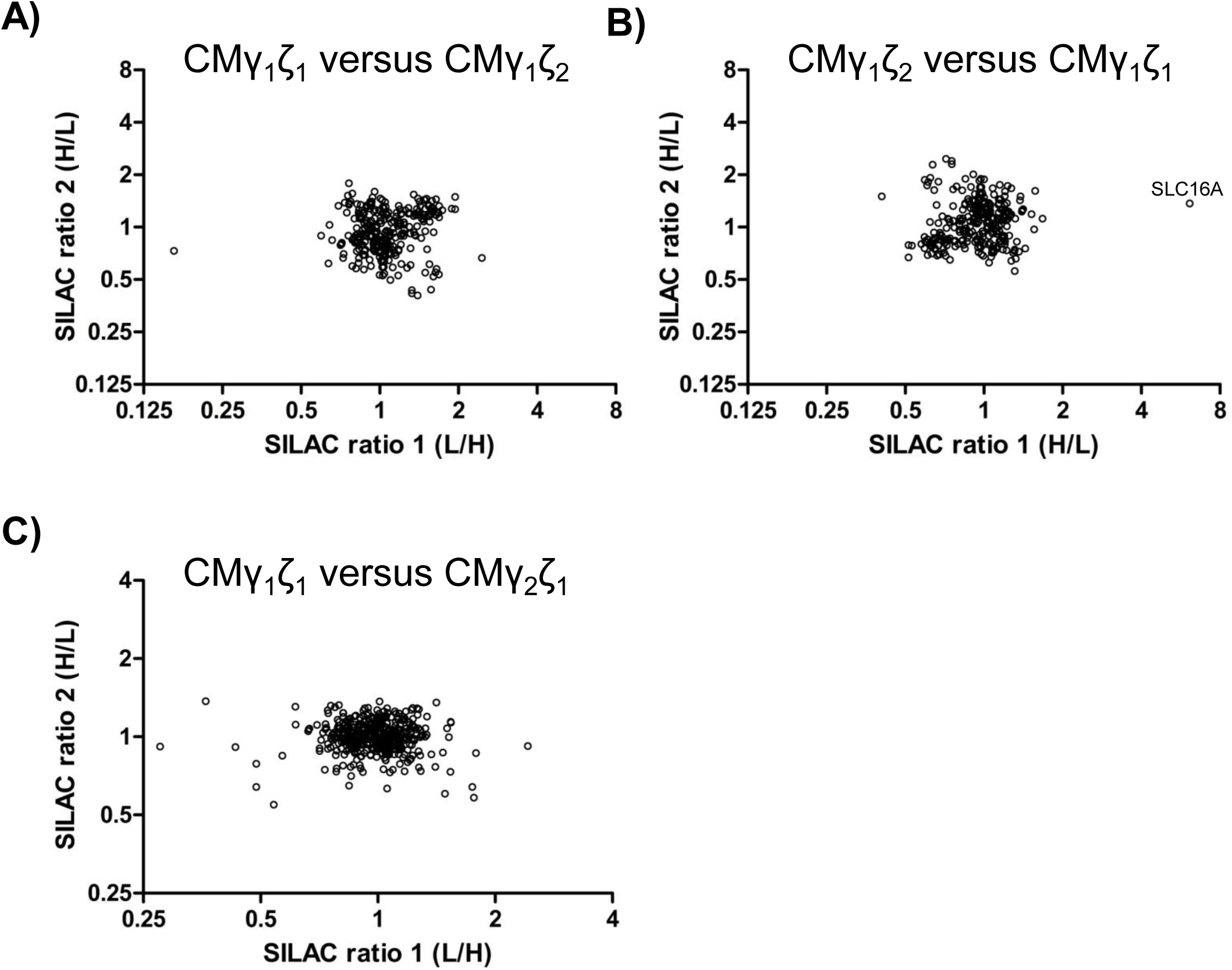
Further comparisons of isotypic COPI vesicles. A-C) Scatter plots representing two independent experiments of a direct comparison of isotypic COPI vesicles indicated. Experiments were performed with switched labels.

**Fig. S3:**
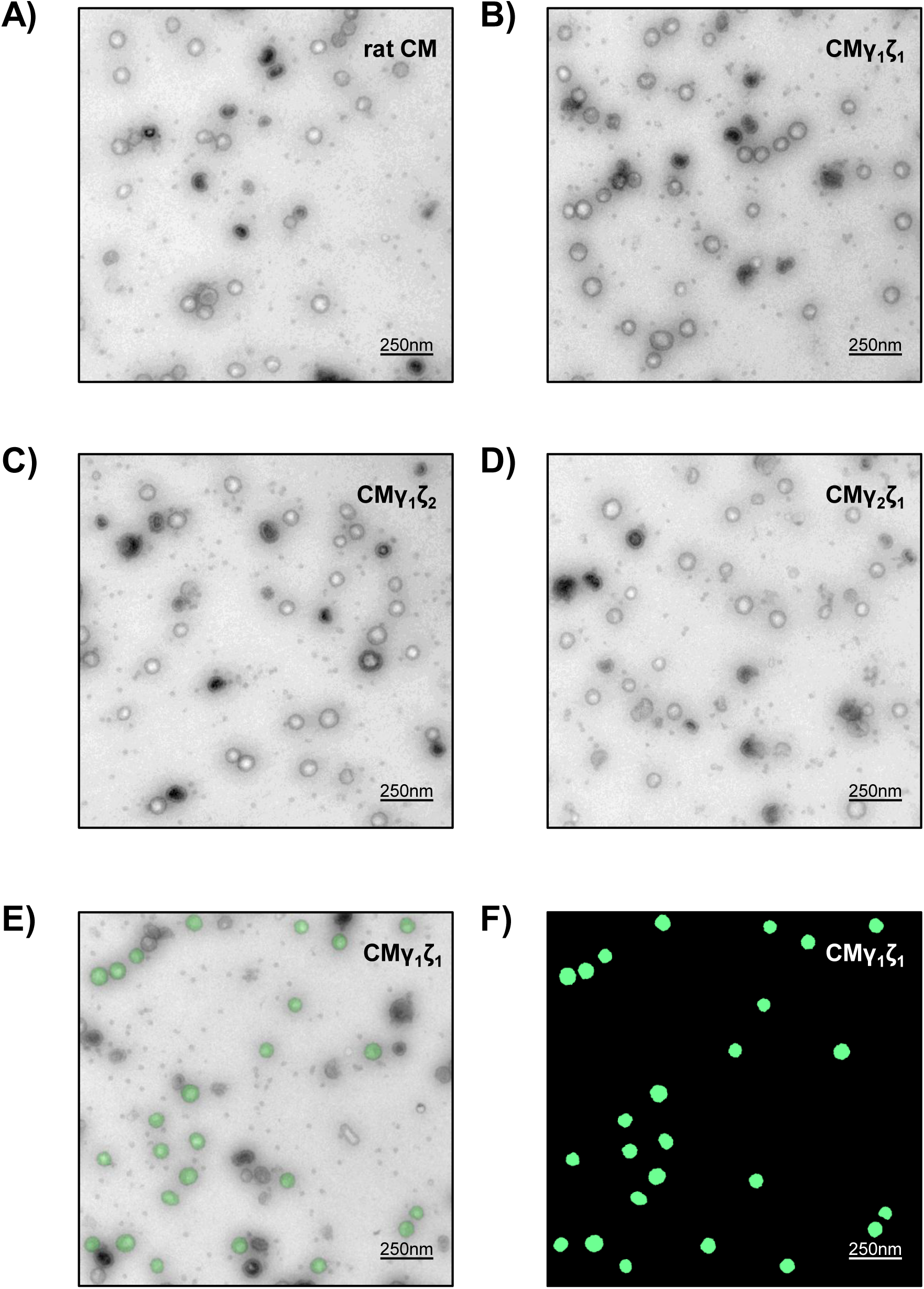
Different coatomer isoforms produce COPI vesicles of similar size. A-D) Representative electron microscopic image of negatively stained COPI vesicles reconstituted with different isoforms of coatomer from rat liver Golgi. E) Determination of vesicle diameters as described in Materials and Methods. Structures taken into account are colored in green. F) Mask of the segmented vesicles shown in E).

